# Fibronectin 1 is a novel biomarker of obstetric antiphospholipid syndrome

**DOI:** 10.1101/2024.01.08.574588

**Authors:** Jinfeng Xu, Tenchi Chung, Zhe Hu, Yuan Tian, Qiao Ling, Xiaodong Wang, Bing Peng

**Affiliations:** Department of Obstetrics and Gynecology, West China Second University Hospital, Sichuan University, Sichuan, China; West China School of Medicine, Sichuan University, Sichuan, China; National key laboratory of biotherapy, Sichuan university, Sichuan, China; The Key Laboratory of Birth Defects and Related Diseases of Women and Children, Sichuan University, Sichuan, China

**Keywords:** Obstetric antiphospholipid syndrome, Fibronectin 1, biomarker

## Abstract

**Objective:** To investigate novel biomarkers and the mechanism of obstetric antiphospholipid syndrome (OAPS).

**Methods:** HTR8/SVneo cells line were treated with plasma from OAPS (OAPS group) and healthy (NC group) pregnant women, respectively. The changes induced by plasma treatment at the transcriptome level were examined by RNA sequencing of the cells. Results were analyzed with bioinformatics tools to elucidate the potential biomarkers. Reverse-transcription quantitative polymerase chain reaction (RT-qPCR), western blotting, hematoxylin and eosin (HE), immunohistochemistry (IHC), and immunofluorescence (IF) were used for subsequent validation.

**Results:** Bioinformatic analysis revealed the expression of Fibronectin 1 (FN1) was significantly increased in OAPS group. On analyzing molecular function, OAPS plasma exposure mainly affected the expression of the genes related to extracellular matrix (ECM) structural constituent. Compared to the NC group, differently expressed genes were mainly annotated to the collagen-containing ECM matrix and the ECM organization. In OAPS group, the protein expression of FN1 was also increased in blood (*p* <0 .05). The mRNA and protein expression of FN1 in placenta tissue were increased (*p* <0 .05) in OAPS group. Massive degeneration and atrophy can be seen in placental villi, with a significant reduction or disappearance of syncytiotrophoblasts and excessive fibrinoid deposition in the villous stroma of OAPS placenta. Both IHC and IF results showed the staining area and intensity of FN1 in the placental villi and stroma were significantly higher in OAPS group.

**Conclusions:** FN1 may play a potential role in the pathogenic mechanisms of OAPS.

**Highlights:** 

## Instruction

Obstetric antiphospholipid syndrome (OAPS) is an autoimmune disorder characterized by adverse pregnancy outcomes with persistent antiphospholipid antibodies (aPLs) positivity ^1, 2^. The incidence of OAPS ranges from 10% to 41.9% of pregnancies in various populations ^3, 4^. According to the Sydney criteria, the diagnosis of APS requires at least one laboratory criterion and one clinical criterion. Laboratory criteria include lupus anticoagulant (LA), or/and anticardiolipin antibody (ACA), or/and anti-beta2 glycoprotein 1 antibody (aβ2GP1) continuous presence. Clinical criteria include: (1) at least three consecutive miscarriages ≤10 gestational weeks, (2) one or more fetal losses (FL) at ≥10 gestational weeks, (3) preterm birth due to eclampsia or severe pre-eclampsia (PE) or placental insufficiency before the 34th week of gestation ^1^. Recently, studies suggested that aPLs-mediated functional damage on placenta plays a role in adverse pregnancy outcomes (APOs) ^5^. The aPLs may interfere with early placentation by reducing trophoblast proliferation and invasiveness ^6^. Moreover, other mechanisms might be involved in the pathogenesis of APS, such as the neutrophil extracellular traps (NETosis)^7^, gut microbiota^8^, and pyroptosis^9^. However, the exact pathogenic mechanisms of OAPS are still poorly understood. Therefore, it’s necessary to investigate novel biomarkers and understand the mechanism of OAPS.

Recently, RNA sequencing (RNA-seq) technology has helped identify pathogenic mechanisms in multiple autoimmune diseases and made it possible to look at the changes at the transcriptomic level and find multiple molecular targets for the experimental treatment at the same time ^10^. Therefore, in the present study, the human HTR8/SVneo cells were treated with plasma from OAPS and healthy pregnant women, respectively. These changes induced by plasma treatment at the transcriptome level were examined by RNA sequencing of the cells. The results were then analyzed with a series of bioinformatics tools to elucidate the potential biomarkers in the cells.

## Materials and Methods

### Ethics statement

The study was approved by the Ethics Committee of West China Second University Hospital, Sichuan University. Written informed consent was obtained from patients before enrolment.

### Patients and Sample collection

This study was performed at West China Second University Hospital, Sichuan University. To explore the pathological mechanism of OAPS, we set up a cohort of OAPS since 1 September 2017, collecting socioeconomic data, pregnancy history, pre-pregnancy condition and comorbidities, aPLs status and medications during pregnancy, maternal comorbidities, fetal and neonatal outcomes. A total of ten OAPS pregnant women were included for sample collection in this study. OAPS was diagnosed based on the 2006 Sydney revised classification criteria. The healthy pregnant women were matched to the cases with respect to age, without autoimmune diseases or other severe internal and surgical disease (Table S1).

Human whole blood from OAPS (OAPS group) and healthy pregnant women (NC group) was collected into tubes ethylenediaminetetraacetic acid (EDTA) to obtain plasma on the day of admission. Term placentas were collected underwent cesarean section from OAPS (n=10) and NC (n=10) group. Samples were flash-frozen in liquid nitrogen and then stored at −80°C for Reverse-transcription quantitative polymerase chain reaction (RT-qPCR) and Western blot. The placental tissues were also fixed with 10% formalin at room temperature. Then, the tissues embedded in paraffin, and cut into 5-μm thick sections for further histological, immunohistochemistry (IHC) and immunofluorescence (IF) staining.

### Cell Culture

The HTR8/SVneo cells were grown in DMEM/F12 supplemented with 10% FBS, 1% penicillin, and 1% streptomycin in 100 mm dishes and maintained at 37°C in a humidified atmosphere containing 5% CO2. The medium was changed every three days. Cells grown to 70%-80% confluence were serum-starved overnight and then treated with 3% OAPS plasma or 3% NC plasma for 48 h.

### RNA Sequencing

Cell samples, treated with 3% OAPS plasma and treated with NC plasma for 48h, were sent to the Beijing Genomics Institute (BGI, Wuhan, China) and sequenced as follows: (1) 1 mg of the total cellular RNA was extracted using the TRIzol® reagent and treated with DNaseI at 37°C for 20 min to digest DNA. Oligo dT magnetic beads were used to select mRNA with polyA tail; (2) the purifified mRNA were fragmented and reverse transcribed to double-strand cDNA (dscDNA) by using the N6 random primer (Sequence: 5’ d(N6) 3’ [N=A, C, G, T]); (3) the dscDNAs were treated by a traditional process, including end repairing with phosphate at the 5′ end and stickiness ‘A’ at the 3′ end, ligation, and adaptor with stickiness ‘T’ at the 3′ end; (4) two specifific primers were used to amplify the ligation product; (5) the double-stranded polymerase chain reaction (PCR) products were heat-denatured to single-strand and circularized by splint oligo and DNA ligase; (6) sequencing was performed on the BGISEQ-500 platform with the prepared library.

### Bioinformatics

Gene expression levels were quantified by using the RSEM software ^11^. DEGseq was adopted to screen the differentially expressed genes (DEGs) between NC and OAPS groups ^12^. The criteria for the selection of the DEGs were fold change ≥ 1.5 and Q-value ≤ 0.05. The volcano plot was performed using the ‘ggplot2’ package in R. The heatmap was constructed using the ‘heatmap.2’ package in R. Analysis of biological process (BP), molecular function (MF), and cellular component (CC) terms was performed using the ToppCluster^13^ and Gene Ontology^14, 15^ databases.

### RNA Isolation and Reverse-Transcription Quantitative Polymerase Chain Reaction (RT-qPCR)

Placental tissues were divided and dissected separately on ice-cooled RNase-free surfaces. Total RNA was extracted using the QuickPrep RNA extraction kit (GE HealthCare). Reverse transcribed using the TaqMan Reverse Transcription Reagent (Applied Biosystems). Quantitative PCR (qPCR) was performed on the CFX Connect Real-Time PCR Detection System using SYBR Green PCR Master Mix (TaKaRa). Primer pairs used were listed in Supplementary (Table S2). Reactions were performed in triplicate under standard thermocycling conditions (one cycle of 94°C for 4 min, followed by 45 cycles of 94°C for 30 s, 58°C for 30 s, and 72°C for 40 s), and the mean threshold cycle number was used. Glyceraldehyde-3-phosphate dehydrogenase (GAPDH) was used as the internal control. The threshold cycle (CT) method (2−ΔΔCT method) was applied.

### Western blot

The total protein of placenta was extracted using RIPA lysis buffer (Beyotime, Beijing, China) with 1% proteinase inhibitor cocktail (Beyotime), and phosphatase inhibitor cocktail (MEC, Houston, USA) followed by centrifugation at 12,000 g and 4°C for 30 min. Protein concentrations of the supernatants were quantified by using the BCA Protein Assay Kit (P0012). The proteins were separated by SDS-PAGE and transferred to polyvinylidene difluoride membranes (Millipore, Temecula, CA) and then blocked in 5% non-fat milk. The blocking step was followed by overnight incubation with primary antibodies, including anti-FN1 (1:1000, 15613-1-AP, Proteintech) and GAPDH (1:5000, 60004-1-Ig, Proteintech) at 4°C. Bands were then washed with 0.1% Tween 20 in tris-buffered saline and incubated with the IgG HRP secondary antibody for 1 hour at room temperature. Then, the bands were detected using the ECL luminol reagent (Merck Millipore, Burlington, MA) and imaged using the ChemiDocTM MP Imaging system (BIO-RAD). The gray value of protein bands was analyzed using the Image J software.

### Histology of placenta

Consecutive 5-μm sections of placentas were produced and processed for H&E staining by a standard protocol. Sections were incubated in Mayer’s Hematoxylin (G1004, Servicebio) for 5 min and briefly washed in tap water. Sections were subsequently differentiated in 70% acid alcohol (70% ethanol and 1% hydrochloric acid) for 10 s, incubated in cold tap water for 30 min, and incubated in eosin (G1002, Servicebio) for 30 s, followed by an increasing ethanol row (70% for 3 min, 95% for 3 min, 100% for 3 min, and 100% for 3 min). After that, the sections were incubated in 100% xylene for 5 min before mounting with Rhamsan gum (WG10004160, Servicebio) and imaged by Research Slide Scanner (OLYMPUS VS200).

### Immunohistochemistry (IHC)

Slides were baked for 1h at 65°C followed by deparaffinization with xylene and a graded series of ethanol dilutions (100%, 85% and 75%) and rehydration, antigen retrieval for 20min at 98°C using the ethylene diamine tetra-acetic acid buffer (pH 8.0) (G1206, Servicebio). Endogenous peroxidase was inhibited by 3% hydrogen peroxide. Blocking by BSA, incubation by primary and secondary antibodies conjugated with HRP were performed sequentially. The primary antibody of FN1 (1:1000, 15613-1-AP, Proteintech) was used. Staining with 3,3-diaminobenzidine tetrahydrochloride substrate solution and counter staining with haematoxylin were followed. The slides were imaged by Research Slide Scanner (OLYMPUS VS200).

### Immunofluorescence (IF)

Placentas were processed in frozen sections (5 μm), fixed by 4% PFA, followed by antigen retrieval for 20min at 98°C using the ethylene diamine tetra-acetic acid buffer (pH 8.0) (G1206, Servicebio) and permeabilization by 0.1% Triton X-100 (w:v) in PBS for 15 min. Slides were blocked in 10% normal donkey serum and then stained with monoclonal anti-FN1 antibody (1:200; 15613-1-AP, Proteintech), followed by Alexa Fluor 488–conjugated goat anti-rabbit antibody (1:500; A11070, Invitrogen, USA). Nuclei were stained with DAPI (40, 6-diamidino-2-phenylindole; 1:5000, Invitrogen). The slides were mounted with a mounting medium (S3023, Dako, Glostrup, Denmark) and observed under a Zeiss Laser scanning confocal microscope (LSM 980, Zeiss, Carl).

### Statistical analysis

All the experimental data were expressed as means ± SD. Two-tailed unpaired Student t tests were conducted using GraphPad Prism 9 (GraphPad Software Inc., USA). A probability value of <0.05 was considered statistically significant.

## Results

### RNA Sequencing

#### RNA-Seq Data Quality

The sequencing generated, on average, 23,923,064 raw sequencing reads and 23,860,042 clean reads after filtering out low-quality reads. The numbers of the reads and their quality metrics for each sample are listed in Supplementary Table S3. The average mapping ratio with the reference genome is 96.13%. The average mapping ratio with the reference genes is 80.06%. A total of 18,044 genes were detected. RSEM-quantifified gene expression levels and the numbers of identifified expressed genes in OAPS and NC groups are shown in Figure S1.

#### Differently Expressed Genes (DEGs)

As shown in Figure 1, there are 715 DEGs between the OAPS and NC group under the criteria of log_2_ fold change ≥ 1.5, and a Q-value <0.05. Compared to the NC group, 520 genes were significantly up-regulated, and 195 were significantly down-regulated in the OAPS group. The hierarchically clustered heatmaps displays that the expression of FN1 was significantly increased in OAPS than that in NC (Figure 2).

**Figure 1.**
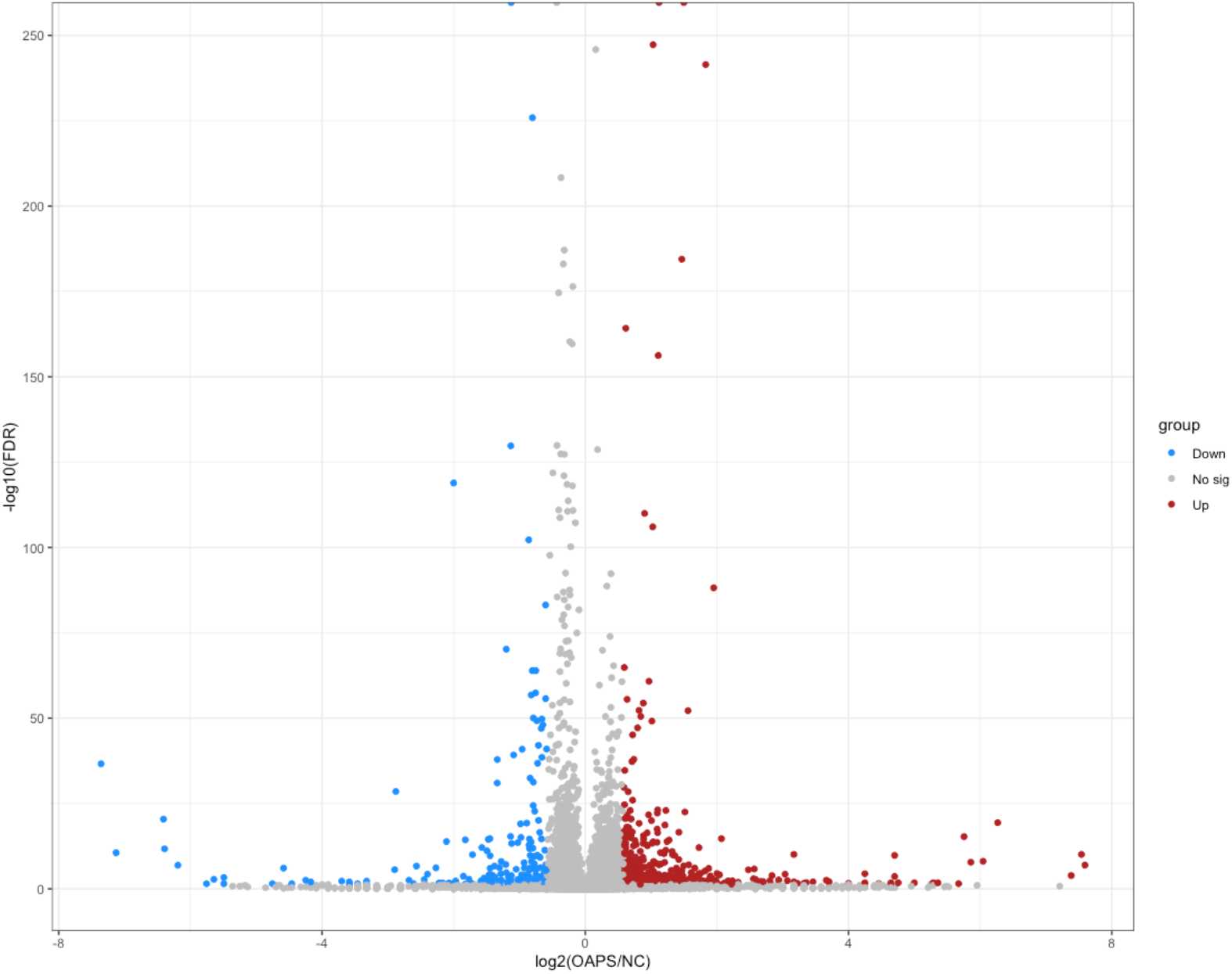
RNA sequencing data. The volcano plot. The X-axis represents the fold change in gene expression between OAPS group and NC group. Y-axis represents the statistical significance of differentially expressed genes. OAPS: obstetric antiphospholipid syndrome group; NC: Healthy Group.

**Figure 2.**
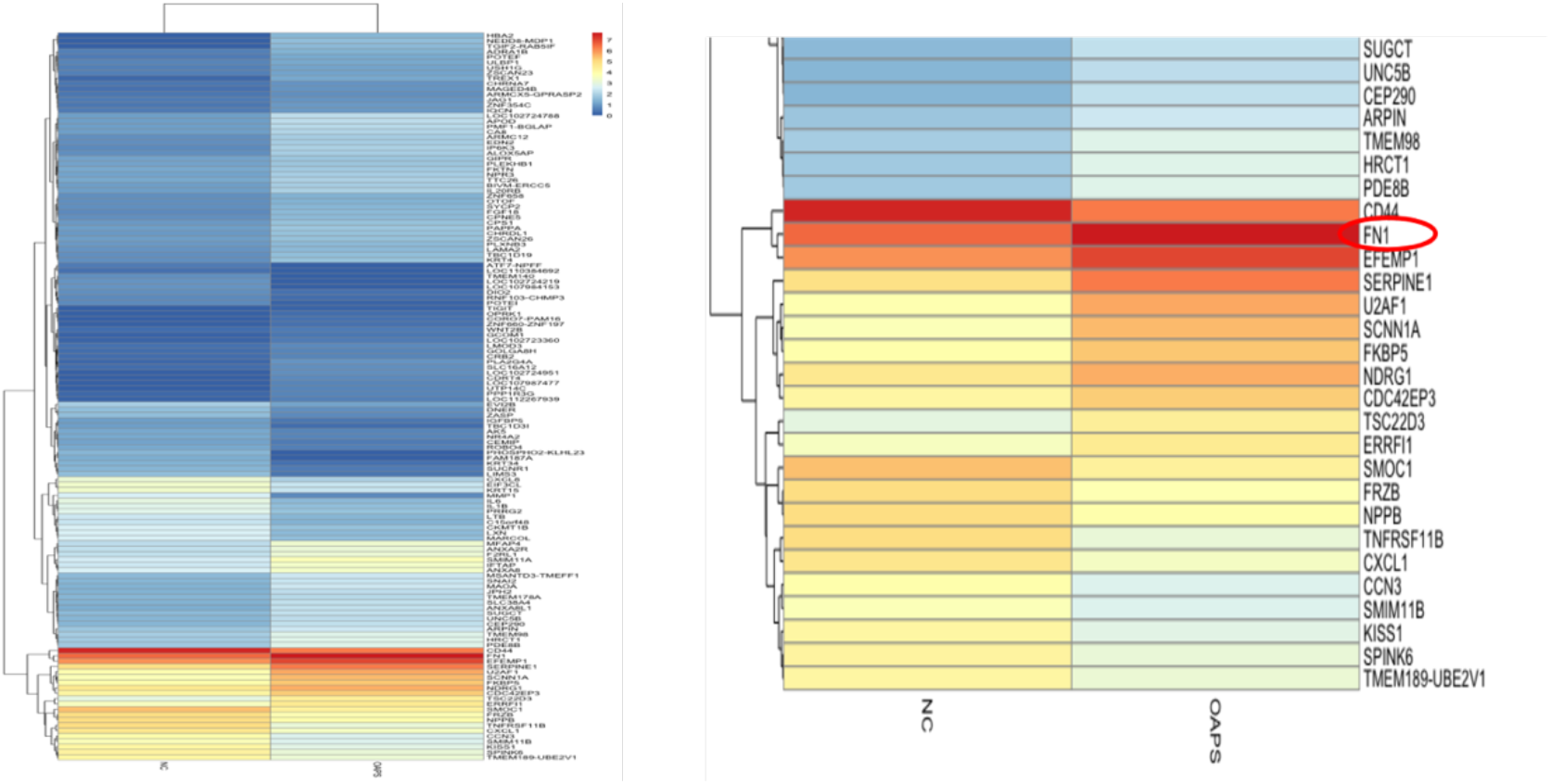
RNA sequencing data. Hierarchical clustering analysis of all the DEGs. The X-axis indicates the sample name. Y-axis indicates the name of identified expressed genes. OAPS: obstetric antiphospholipid syndrome group; NC: Healthy Group.

#### Gene Ontology (GO) Analysis

Based on the GO annotations of the DEGs after OAPS and NC plasma treatment, we had a better understanding of the effect of OAPS plasma on HTR8/SVneo cells in terms of the three GO classififications-molecular function (MF), cellular component (CC), and biological process (BP). On analyzing MF, Figure 3 shows that, compared to NC, OAPS plasma exposure mainly affected the expression of the genes related to extracellular matrix (ECM) structural constituent. Similarly, CC analysis indicates that DEGs were mainly annotated to the collagen-containing ECM (Figure 3). With regards to BP, compared to NC, DEGs were also related to the ECM organization after OAPS plasma exposure (Figure 3).

**Figure 3.**
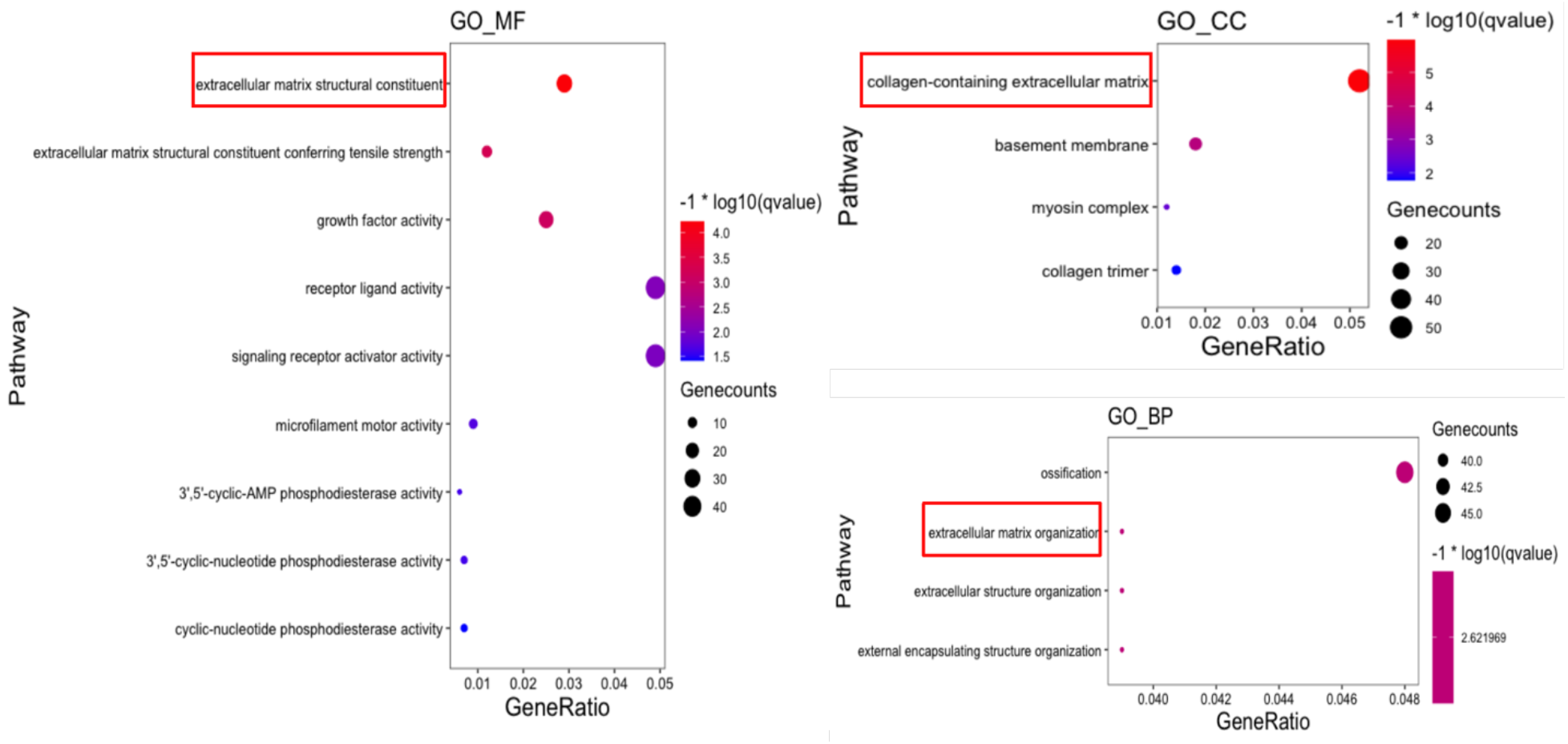
RNA sequencing data. Significantly different GO terms between the two groups. The expression level of the genes in two groups and the resulting differentially expressed genes (DEGs) were subjected to the gene ontology (GO) analysis in terms of three GO classification-molecular function (MF), cellular component (CC), and biological process (BP).

### RT-qPCR and Western blot

The expression level of FN1 mRNA of placenta tissue was increased in OAPS group compared with that in NC group (*p* <0 .05; Figure 4(left)). Like the results of mRNA, the protein expression of FN1 in placenta showed an increase in OAPS group compared with NC group. We didn’t detect FN1 mRNA expression in blood both in OAPS group and NC group (*p* <0 .05; Figure 4(right)). However, the protein expression of FN1 of OAPS group was increased in blood compared with NC group (*p* <0 .05; Figure 4(right)).

**Figure 4.**
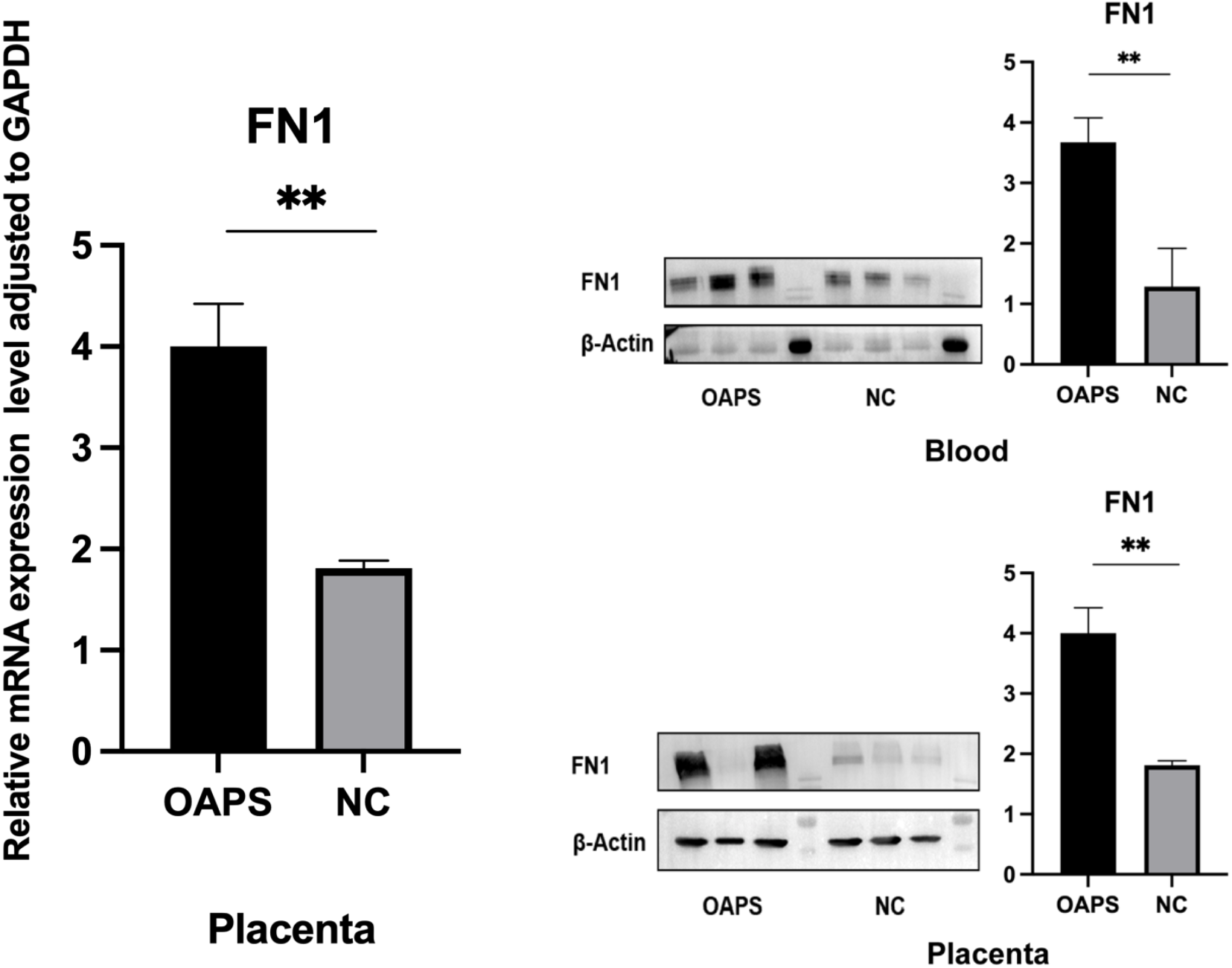
Relative mRNA expression of FN1 in placenta were examined by RT-PCR in OAPS and NC groups (left). The protein of FN1 both in blood and placenta by western blot PCR in OAPS and NC groups (right). β-Actin was used as the internal standard. **p <0 .05 vs aPL group. OAPS: obstetric antiphospholipid syndrome group; NC: Healthy Group.

### Placental histological analysis, IHC and IF

Massive degeneration and atrophy can be seen in placental villi, with a significant reduction or disappearance of syncytiotrophoblasts and excessive fibrinoid deposition in the villous stroma of OAPS placenta. The basal membrane of the capillary walls showed slight thickening. (Figure 5(left)).

**Figure 5.**
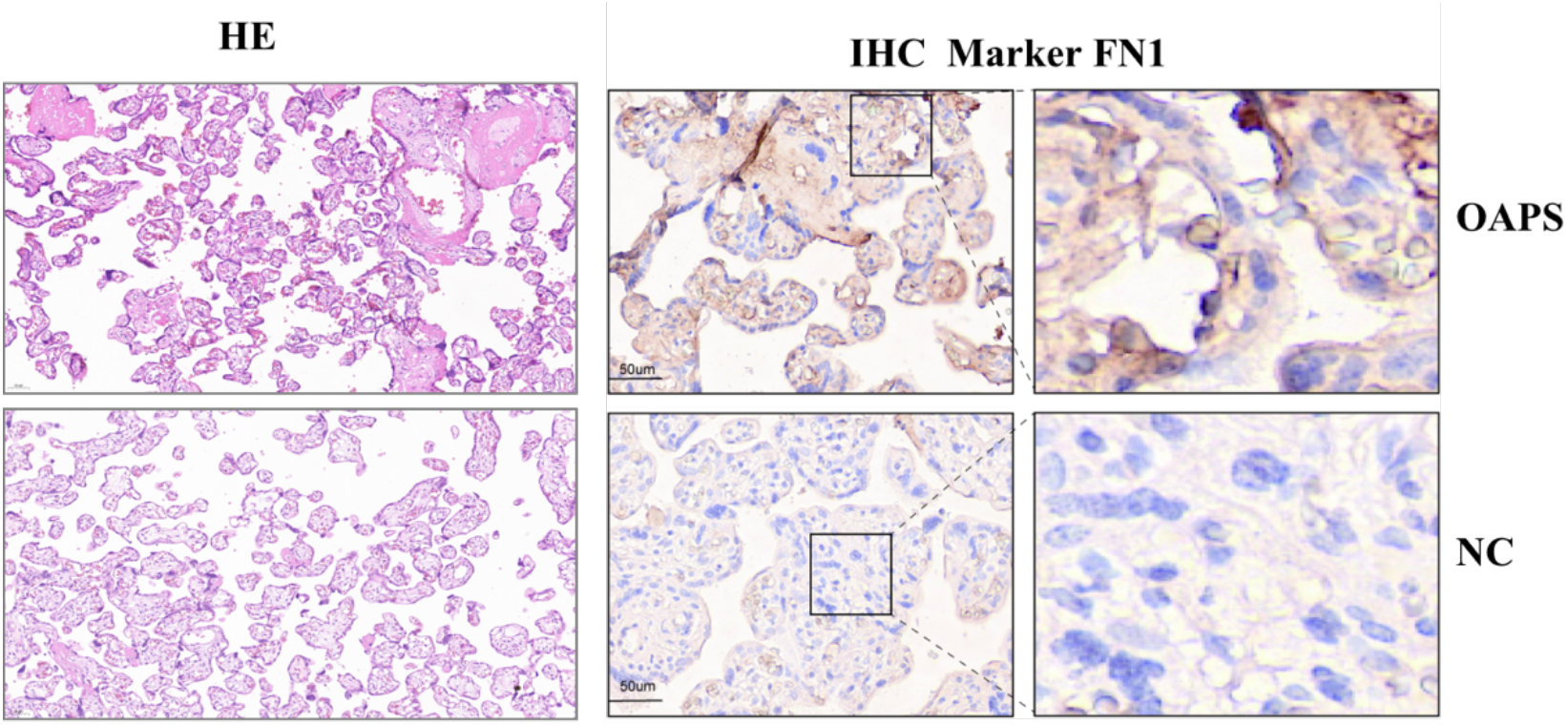
Histological analysis. (A) H&E Placental tissues specimens were stained with H&E staining. Original magnification 20×. The cases were listed as OAPS, and NC. (B) Immunohistochemistry (IHC) analysis. Representative images of IHC of proteins (FN1) expression from OAPS and normal tissues. IHC images were taken under 20×. OAPS: obstetric antiphospholipid syndrome group; NC: Healthy Group.

IHC was performed to detect the expression levels of FN1 in the maternal-fetal interface placental tissues (Figure 5(right)). FN1 was mainly localized on the trophoblast surface, cytoplasm, basal membrane, and ECM. In OAPS group, the staining area and intensity of FN1 in the placental villi and stroma were significantly higher compared to NC group. IF staining of placental tissues also revealed a significant up-regulation of FN1 expression in OAPS group when compared to NC group (Figure 6).

**Figure 6.**
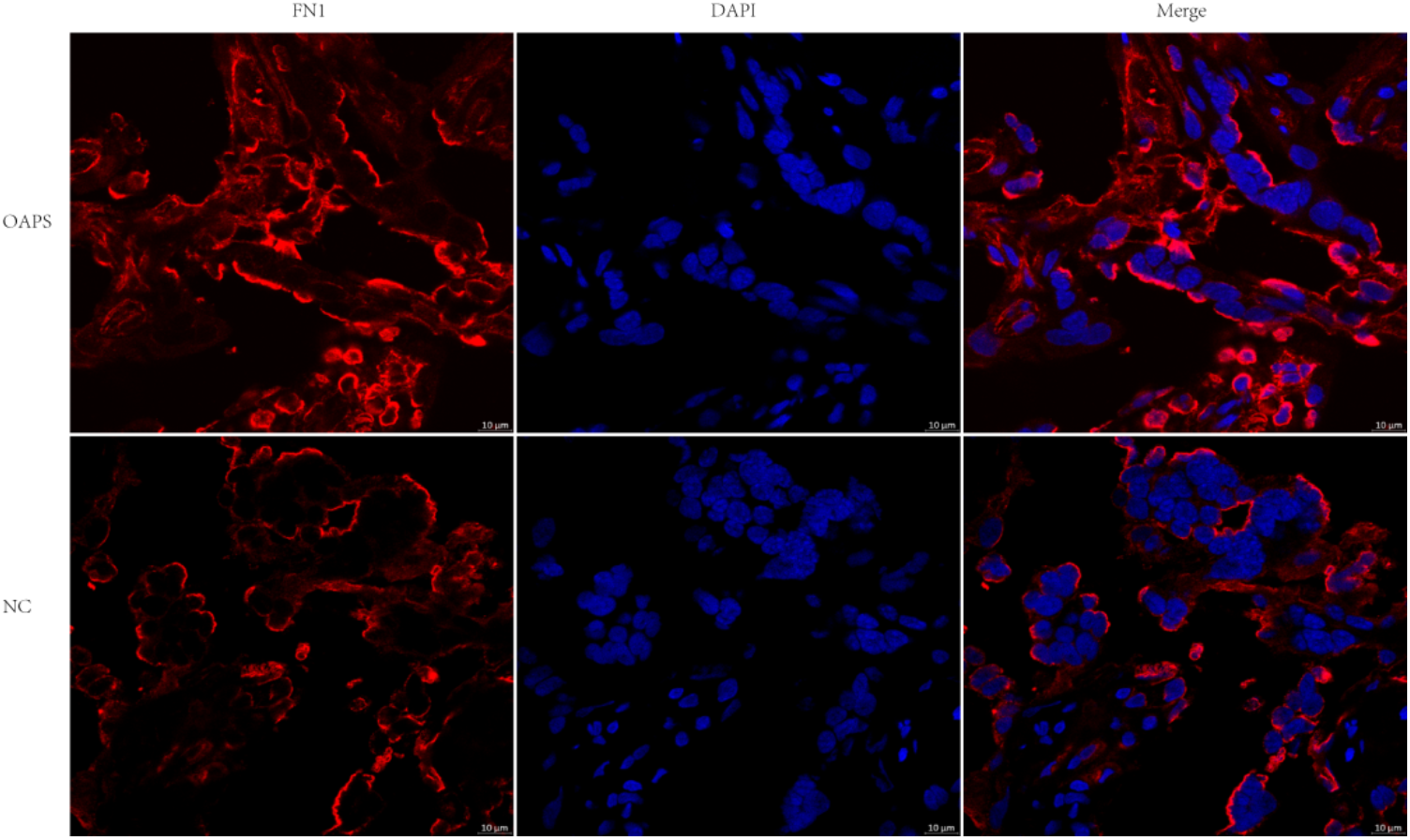
Immunofluorescence staining of FN1 (red) in human placenta samples. Nuclei were counterstained with DAPI (blue). Scale bars, 10 μm. The cases were listed as OAPS, and NC. OAPS: obstetric antiphospholipid syndrome group; NC: Healthy Group.

## Discussion

Placenta is crucial for fetal growth during pregnancy, exchanging nutrients and wastes between mother and fetus. As reported, the disorder of maternal-fetal interface plays an important role in aPLs-related APOs ^5^. Pathological features, like defective endovascular extravillous trophoblast (enEVT) invasion, impaired spiral artery remodeling, villi calcification, and decreased vasculosyncytial membranes, are common in maternal-fetal interface of OAPS pregnant women, indeed ^5, 16^.

In our study, FN1 was significantly expressed in OAPS placental tissues than those in NC placental tissues. Generally, FN1 is present in the ECM, which plays a key role in cell adhesion, growth, migration, differentiation, and maintaining cell morphology ^17^. ECM homeostasis is crucial for placenta development and function. Trophoblasts express integrins, E-cadherin, and proteases, degrading ECM proteins such as FN1 to promote migration and invasion^18^. Normally, as the pregnancy progresses, the amount of ECM decreases while the number of vessels increase, supporting fetus growth.^19, 20^ However, once this homeostasis is perturbed, either ECM production increased or ECM degradation decreased, excessive ECM deposition can contribute to pathological conditions.

Our study found that excessive FN1 deposition in the villous stroma in OAPS group. On the one hand, excessive FN1 deposition may cause shallow invasion of EVT and inadequate remodeling of spiral arteries, both of which are features of PE. At the same time, nutrients and gas exchange decreased, leading to fetal growth restriction. On the other hand, excessive FN1 accumulation is a feature of fibrosis^21^, such as kidney fibrosis and liver fibrosis, causing increased tissue stiffness and progressive organ dysfunction^22^. Similarly, our results showed that excessive FN1 deposition may be associated with placenta fibrosis, thereby leading to reduced blood flow from uterine to fetus. At early stage of placenta fibrosis, the placenta can compensate. That’s the reason why some pregnant women of OAPS escape spontaneous abortion at first trimester. Unfortunately, with the fibrosis exacerbation at mid-to-late term, the remaining normal placental tissue can’t exert proper function, resulting in devastating complication-intrauterine demise. Excessive FN1 deposition may result from the dysfunction of trophoblasts. Recently, Chunwei Cao et.al reported that aPLs might bind to β2GPI complex expressed by trophoblasts, leading to FN1 deposition ^23^. In agreement with them, our RNA-Seq results indicated that the expression of FN1 was significantly increased in HTR8/SVneo cells treated by OAPS plasma. Moreover, IHC and IF results demonstrated that the intensity of FN1 was significantly higher in OAPS group, which was mainly localized on the trophoblast surface, cytoplasm, basal membrane, and ECM.

Our study had several limitations. Firstly, the sample size for RNA-Seq should be increased in the future. Next, excessive FN1 deposition was the results of the interaction of aPLs and trophoblasts. Thus, further studies, especially single cell RNA-Seq of OAPS placenta, are needed to investigate the potential mechanism of OAPS. Also, experiments in vivo are needed to validate the potential mechanism.

## Conclusions

FN1 may play a potential role in the pathogenic mechanisms of OAPS. More studies in vitro and in vivo are needed to elucidate the pathogenesis of OAPS.

## Notes

### Competing Interest Statement

The authors have declared no competing interest.

## Reference

1. Miyakis S, Lockshin MD, Atsumi T, et al. International consensus statement on an update of the classification criteria for definite antiphospholipid syndrome (APS). J Thromb Haemost 2006; 4: 295–306. DOI: 10.1111/j.1538-7836.2006.01753.x.

2. Esteve-Valverde E, Ferrer-Oliveras R and Alijotas-Reig J. Obstetric antiphospholipid syndrome. Rev Clin Esp (Barc) 2016; 216: 135–145. 20151023. DOI: 10.1016/j.rce.2015.09.003.

3. Jiang H, Wang CH, Jiang N, et al. Clinical characteristics and prognosis of patients with isolated thrombotic vs. obstetric antiphospholipid syndrome: a prospective cohort study. Arthritis Res Ther 2021; 23: 138. 20210508. DOI: 10.1186/s13075-021-02515-w.

4. Ruiz-Irastorza G, Crowther M, Branch W, et al. Antiphospholipid syndrome. Lancet 2010; 376: 1498–1509. 20100906. DOI: 10.1016/s0140-6736(10)60709-x.

5. Viall CA and Chamley LW. Histopathology in the placentae of women with antiphospholipid antibodies: A systematic review of the literature. Autoimmun Rev 2015; 14: 446–471. 20150122. DOI: 10.1016/j.autrev.2015.01.008.

6. Spinillo A, Bellingeri C, Cavagnoli C, et al. Maternal and foetal placental vascular malperfusion in pregnancies with anti-phospholipid antibodies. Rheumatology (Oxford) 2021; 60: 1148–1157. DOI: 10.1093/rheumatology/keaa499.

7. Lu Y, Dong Y, Zhang Y, et al. Antiphospholipid antibody-activated NETs exacerbate trophoblast and endothelial cell injury in obstetric antiphospholipid syndrome. J Cell Mol Med 2020; 24: 6690–6703. 20200505. DOI: 10.1111/jcmm.15321.

8. Ruff WE, Vieira SM and Kriegel MA. The role of the gut microbiota in the pathogenesis of antiphospholipid syndrome. Curr Rheumatol Rep 2015; 17: 472. DOI: 10.1007/s11926-014-0472-1.

9. Di L, Zha C and Liu Y. Platelet-derived microparticles stimulated by anti-β(2)GPI/β(2)GPI complexes induce pyroptosis of endothelial cells in antiphospholipid syndrome. Platelets 2023; 34: 2156492. DOI: 10.1080/09537104.2022.2156492.

10. Zeng L, Yang K, Zhang T, et al. Research progress of single-cell transcriptome sequencing in autoimmune diseases and autoinflammatory disease: A review. J Autoimmun 2022; 133: 102919. 20221012. DOI: 10.1016/j.jaut.2022.102919.

11. Li B and Dewey CN. RSEM: accurate transcript quantification from RNA-Seq data with or without a reference genome. BMC Bioinformatics 2011; 12: 323. 20110804. DOI: 10.1186/1471-2105-12-323.

12. Wang L, Feng Z, Wang X, et al. DEGseq: an R package for identifying differentially expressed genes from RNA-seq data. Bioinformatics 2010; 26: 136–138. 20091024. DOI: 10.1093/bioinformatics/btp612.

13. Kaimal V, Bardes EE, Tabar SC, et al. ToppCluster: a multiple gene list feature analyzer for comparative enrichment clustering and network-based dissection of biological systems. Nucleic Acids Res 2010; 38: W96–102. 20100519. DOI: 10.1093/nar/gkq418.

14. Ashburner M, Ball CA, Blake JA, et al. Gene ontology: tool for the unification of biology. The Gene Ontology Consortium. Nat Genet 2000; 25: 25–29. DOI: 10.1038/75556.

15. The Gene Ontology Resource: 20 years and still GOing strong. Nucleic Acids Res 2019; 47: D330–d338. DOI: 10.1093/nar/gky1055.

16. Sebire NJ, Fox H, Backos M, et al. Defective endovascular trophoblast invasion in primary antiphospholipid antibody syndrome-associated early pregnancy failure. Hum Reprod 2002; 17: 1067–1071. DOI: 10.1093/humrep/17.4.1067.

17. Ji J, Chen L, Zhuang Y, et al. Fibronectin 1 inhibits the apoptosis of human trophoblasts by activating the PI3K/Akt signaling pathway. Int J Mol Med 2020; 46: 1908–1922. 20200923. DOI: 10.3892/ijmm.2020.4735.

18. Rosario GX, Ain R, Konno T, et al. Intrauterine fate of invasive trophoblast cells. Placenta 2009; 30: 457–463. 20090402. DOI: 10.1016/j.placenta.2009.02.008.

19. Zhou Y, Fisher SJ, Janatpour M, et al. Human cytotrophoblasts adopt a vascular phenotype as they differentiate. A strategy for successful endovascular invasion? J Clin Invest 1997; 99: 2139–2151. DOI: 10.1172/jci119387.

20. Kaufmann P, Black S and Huppertz B. Endovascular trophoblast invasion: implications for the pathogenesis of intrauterine growth retardation and preeclampsia. Biol Reprod 2003; 69: 1–7. 20030305. DOI: 10.1095/biolreprod.102.014977.

21. Pakshir P and Hinz B. The big five in fibrosis: Macrophages, myofibroblasts, matrix, mechanics, and miscommunication. Matrix Biol 2018; 68-69: 81–93. 20180131. DOI: 10.1016/j.matbio.2018.01.019.

22. Cox TR and Erler JT. Remodeling and homeostasis of the extracellular matrix: implications for fibrotic diseases and cancer. Dis Model Mech 2011; 4: 165–178. 20110214. DOI: 10.1242/dmm.004077.

23. Cao C, Bai S, Zhang J, et al. Understanding recurrent pregnancy loss: recent advances on its etiology, clinical diagnosis, and management. Med Rev (Berl) 2022; 2: 570–589. 20221219. DOI: 10.1515/mr-2022-0030.

