## Supplemental Figure 1 for "Fibronectin 1 is a novel biomarker of obstetric antiphospholipid syndrome"

**Supplementary Figure S1**

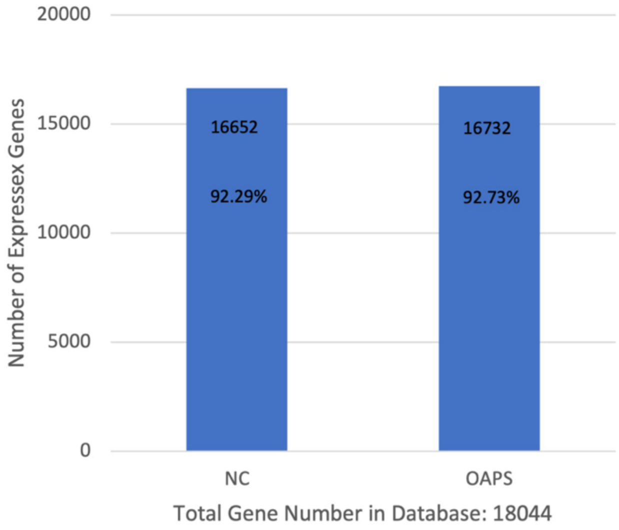


Figure S1 | RNA sequencing data. The number of identified genes. The X-axis indicates the sample name. Y-axis indicates the number of identified expressed genes. The proportion at the top of each bar equals the expressed gene number divided by the total gene number reported in the database. NC: Control Group; OAPS: obstetric antiphospholipid syndrome.
