## Supplemental Table 1 for "Fibronectin 1 is a novel biomarker of obstetric antiphospholipid syndrome"

**Supplementary Table S1.** Characteristics of pregnant women enrolled in this study.

| **Characteristic** | **OAPS (N=10)** | **NC (N=10)** |
|---|---|---|
| Age (y) | 31.4 | 31.2 |
| Gestational weeks | 12.7 | 12.6 |
| Clinical criteria | | |
| RSA | 2 | 0 |
| Late fetal loss | 8 | 0 |
| Laboratory criteria | | |
| LA | 7 | 0 |
| ACA IgG/IgM | 2 | 0 |
| aβ2GP1 IgG/IgM | 2 | 0 |

Notes: N: number; OAPS: pregnant women with obstetric antiphospholipid syndrome; NC: healthy pregnant women; RSA: recurrent spontaneous abortion; LA: lupus anticoagulant; ACA: anticardiolipin antibody; aβ2GP1: anti-beta2 glycoprotein 1 antibody.
