## Supplemental Table 3 for "Fibronectin 1 is a novel biomarker of obstetric antiphospholipid syndrome"

**Supplementary Table S3.** The numbers of the reads and their quality metrics for each sample.

| Sample | Total Raw Reads (Mb) | Total Clean Reads (Mb) | Total Clean Bases (Gb) | Clean Reads Q20 (%) | Clean Reads Q30 (%) | Clean Reads Ratio (%) |
|---|---|---|---|---|---|---|
| NC | 23.92 | 23.86 | 1.19 | 98.25 | 94.55 | 99.72 |
| OAPS | 23.92 | 23.86 | 1.19 | 98.32 | 94.74 | 99.75 |

Notes: Samples: Sample names; Total Raw Reads (Mb): The reads amount before filtering, Unit: Mb; Total Clean Reads (Mb): The reads amount after filtering, Unit: Mb; Total Clean Bases (Gb): The total base amount after filtering, Unit: Gb; Clean Reads Q20 (%): The Q20 value for the clean reads; Clean Reads Q30 (%): The Q30 value for the clean reads; Clean Reads Ratio (%): The ratio of the amount of clean reads; OAPS: pregnant women with obstetric antiphospholipid syndrome; NC: healthy pregnant women.
